# LGR6 is necessary for attaining peak bone mass and regulates osteogenesis through differential ligand use

**DOI:** 10.1101/2021.07.16.452698

**Authors:** Vikram Khedgikar, Julia F. Charles, Jessica A. Lehoczky

## Abstract

Leucine-rich repeat containing G-protein-coupled receptor 6 (LGR6) is a marker of osteoprogenitor cells and is dynamically expressed during in vitro osteodifferentation of mouse and human mesenchymal stem cells (MSCs). While the Lgr6 genomic locus has been associated with osteoporosis in human cohorts, the precise molecular function of LGR6 in osteogenesis and maintenance of bone mass are not yet known. In this study, we performed in vitro Lgr6 knockdown and overexpression experiments in murine osteoblastic cells and find decreased Lgr6 levels results in reduced osteoblast proliferation, differentiation, and mineralization. Consistent with these data, overexpression of Lgr6 in these cells leads to significantly increased proliferation and osteodifferentiation. To determine whether these findings are recapitulated in vivo, we performed microCT and ex vivo osteodifferentiation analyses using our newly generated CRISPR-Cas9 mediated Lgr6 mouse knockout allele (Lgr6-KO). We find that ex vivo osteodifferentiation of Lgr6-KO primary MSCs is significantly reduced, and 8 week-old Lgr6-KO mice have less trabecular bone mass as compared to Lgr6 wildtype controls, indicating that Lgr6 is necessary for normal osteogenesis and to attain peak bone mass. Toward mechanism, we analyzed in vitro signaling in the context of two LGR6 ligands, RSPO2 and MaR1. We find that RSPO2 stimulates LGR6-mediated WNT/β-catenin signaling whereas MaR1 stimulates LGR6-mediated cAMP activity, suggesting two ligand-dependent functions for LGR6 receptor signaling during osteogenesis. Collectively, this study reveals that Lgr6 is necessary for wildtype levels of proliferation and differentiation of osteoblasts, and achieving peak bone mass.

## INTRODUCTION

Bone is a dynamic tissue that is continuously remodeled throughout life to maintain its strength and structural integrity. This remodeling relies on osteoclast mediated resorption of old or damaged bone and generation of new bone by osteoblasts. Maintenance of bone homeostasis requires a precise balance of these processes, whereby excess resorption or reduced bone formation can result in bone diseases such as osteoporosis and associated fragility fractures^1^. It is estimate that over 10 million people in the U.S. have osteoporosis, a diagnosis that is heavily skewed towards post-menopausal women^2^. This diagnosis has a strong correlation with fragility fractures; as many as half of women over the age of 50 are at risk for age-related bone fractures^2,3^. Antiresorptive drugs can stabilize bone mass and reduce fracture risk in osteoporosis patients^4^. While clinically effective, these drugs have drawbacks including side effects, discontinuation rebound resorption, and ineffectiveness in directly stimulating new bone growth^4–6^. Existing anabolic therapies can be successful in stimulating new bone growth; however, their effects decrease over time and gains in bone mass are lost after discontinuation^7,8^. Off target effects including cardiovascular complications and the potential for cancer are also a concern^9–11^. It follows that there is a significant need to develop new anabolic therapies, which requires identification of new targets and increased understanding of the molecular signaling in osteogenesis. Leucine-rich repeat containing G-protein-coupled receptor 6 (LGR6) as described in this report represents a new molecular target toward this goal. LGR6 is a 7-transmembrane G-protein-coupled receptor that has been identified as a marker of adult progenitor cell populations in diverse tissues including skin, nail, lung, and bone^12–15^. Previously, we demonstrated that Lgr6 is a marker of osteoprogenitors in mice and is dynamically expressed during in vitro osteodifferentiation of mesenchymal stem cells (MSCs), suggesting an osteogenic function^13,14^. These findings translate to human osteogenesis in that Lgr6 is also dynamically expressed during in vitro osteodifferentiation of human MSCs and osteoblasts^16,17^. Importantly, existing data support a function for LGR6 in bone homeostasis and repair; Lgr6 is necessary for mouse digit bone regeneration, and targeted gene sequencing of postmenopausal Chinese women found Lgr6 to be associated with osteoporosis^13,18^.

There are two defined ligands for the LGR6 receptor: R-spondin proteins (RSPOs: RSPO1, RSPO2, RSPO3, and RSPO4) and the lipid metabolite maresin-1 (MaR1), though these ligand-receptor interactions have not yet been explicitly defined in bone^19,20^. RSPO bound LGR6 agonizes WNT signaling, and while all four RSPOs can potentiate this signaling, in vitro studies find RSPO2 binds LGR6 with the highest affinity^19^. The structural interaction and downstream signaling has not yet been described specifically in bone, but data from each protein alone support the putative interaction. Overexpression of Lgr6 in MC3T3-E1 cells increases WNT/β-catenin signaling and differentiation of osteoblasts^21^. Similarly, RSPO2 has an anabolic effect on osteoblasts in vitro, and Ocn-cre;Rspo2-flox conditional knockout mice have reduced osteogenesis and bone mass^22,23^. Beyond WNT signaling, MaR1 bound LGR6 potentiates G-protein mediated signaling and cAMP activity as was recently defined in the context of immunoresolution in human and mouse phagocytes^20^. This ligand-receptor signaling interaction has not yet been reported in other cell types, but MaR1 has been associated with anabolic bone phenotypes. Using an aged mouse fracture model, MaR1 treatment resulted in improved bone regeneration, and in a separate study, local application of MaR1 to rat extracted molar sockets led to accelerated wound healing and alveolar bone regeneration^24,25^. Broadly speaking, WNT signaling and G-protein/cAMP signaling are essential and well-studied in the context of osteogenesis^26,27^. We hypothesize that LGR6 is a unique receptor in bone, capable of regulating both signaling pathways in response to differential ligand use.

In this report, we build upon the previous data and use in vitro knockdown and overexpression of Lgr6 in immortalized and primary mouse MSCs and calvarial osteoblasts to provide further evidence that Lgr6 regulates proliferation, differentiation, and mineralization of osteoblasts. By immunohistochemistry, we detect LGR6 expressing cells within the bone marrow as well as on the surface of the femoral trabeculae. Genetic lineage analysis of these cells reveals they contribute to the osteoblasts of the trabecular surface and the adjacent cortical bone. Using CRISPR-Cas9, we engineered a new Lgr6 knockout allele. MicroCT analysis of mutant and wildtype littermates reveal Lgr6 knockout mice have significantly reduced trabecular bone mass. Towards mechanism, we use in vitro signaling assays to show that RSPO2 stimulates LGR6-mediated WNT/β-catenin signaling and MaR1 stimulates LGR6-mediated cAMP activity, demonstrating differential ligand-induced signaling during osteogenesis. Taken together, this study anchors Lgr6 to osteogenesis and lays the foundation for a novel inroad to anabolic therapies.

## RESULTS

### Lgr6 regulates MSC proliferation and osteodifferentiation

Previously, we established Lgr6 as a marker of osteoprogenitors with dynamic expression during osteogenesis in mice^14^. In line with this, a recent report ties Lgr6 expression to WNT/β-catenin signaling and in vitro differentiation of MC3T3E1 immortalized calvarial cells, though does not explore signaling or function in bone marrow derived MSCs, primary osteoblasts, or in vivo^21^. Building on these data, to determine if expression of Lgr6 contributes to the proliferation or differentiation in primary osteoprogenitors, we used mouse bone marrow derived MSCs. We first knocked-down endogenous Lgr6 expression using an in vitro siRNA-mediated approach (Supplemental Figure 1). 24 hours post-transfection, we found an 11% reduction in proliferation of Lgr6 knockdown cells as compared to control siRNA knockdown cells by WST-1 based cell proliferation assays (p=0.011) (Figure 1a). To determine if Lgr6 is necessary for osteoblastogenesis in primary MSCs, we used Lgr6 siRNA-mediated knockdown during in vitro osteodifferentiation and assessed differentiation markers and mineralization. Runx2 expression was significantly decreased 1 and 3 days post-osteoinduction in Lgr6 knockdown cells compared to controls (day 1: 46%, p=8.53E-05; day 3: 36%, p=0.002) (Figure 1b). Similarly, Sp7 expression in Lgr6 knockdown cells was decreased as compared to controls (day 1: 56%, p=0.030; day 3: 72%, p=0.002) (Figure 1c). We found alkaline phosphatase (ALP) activity was 19% reduced by 7 days post-osteoinduction in Lgr6 knockdown cells as compared to negative siRNA controls (p=0.0016) (Figure 1d). Consistent with this finding, mineralization of Lgr6 knockdown cells was reduced by 16% (p=0.016) as determined by quantification of alizarin red staining (ARS) 18 days post-osteoinduction (Figure 1e). We performed separate Lgr6 siRNA knockdown experiments in primary mouse calvarial osteoblasts (MCO) to validate our MSC data. As with MSCs, Lgr6 knockdown MCOs showed reduced proliferation by WST-1 (10%, p=0.019) and decreased expression of Runx2 (60%, p=0.0006) and Sp7 (38%, p=4.66E-05) 2 days post-induction (Supplemental Figure 2a-c). Also consistent with the MSC data, Lgr6 knockdown MCOs had reduced ALP activity (15%, p=0.005) and ARS staining (18%, p=0.006) at 7 and 21 days post-induction, respectively (Supplemental Figure 2d-e). Collectively, these data demonstrate that Lgr6 is necessary for normal levels of proliferation and osteodifferentiation of mouse primary MSCs and MCOs.

**Figure 1.**
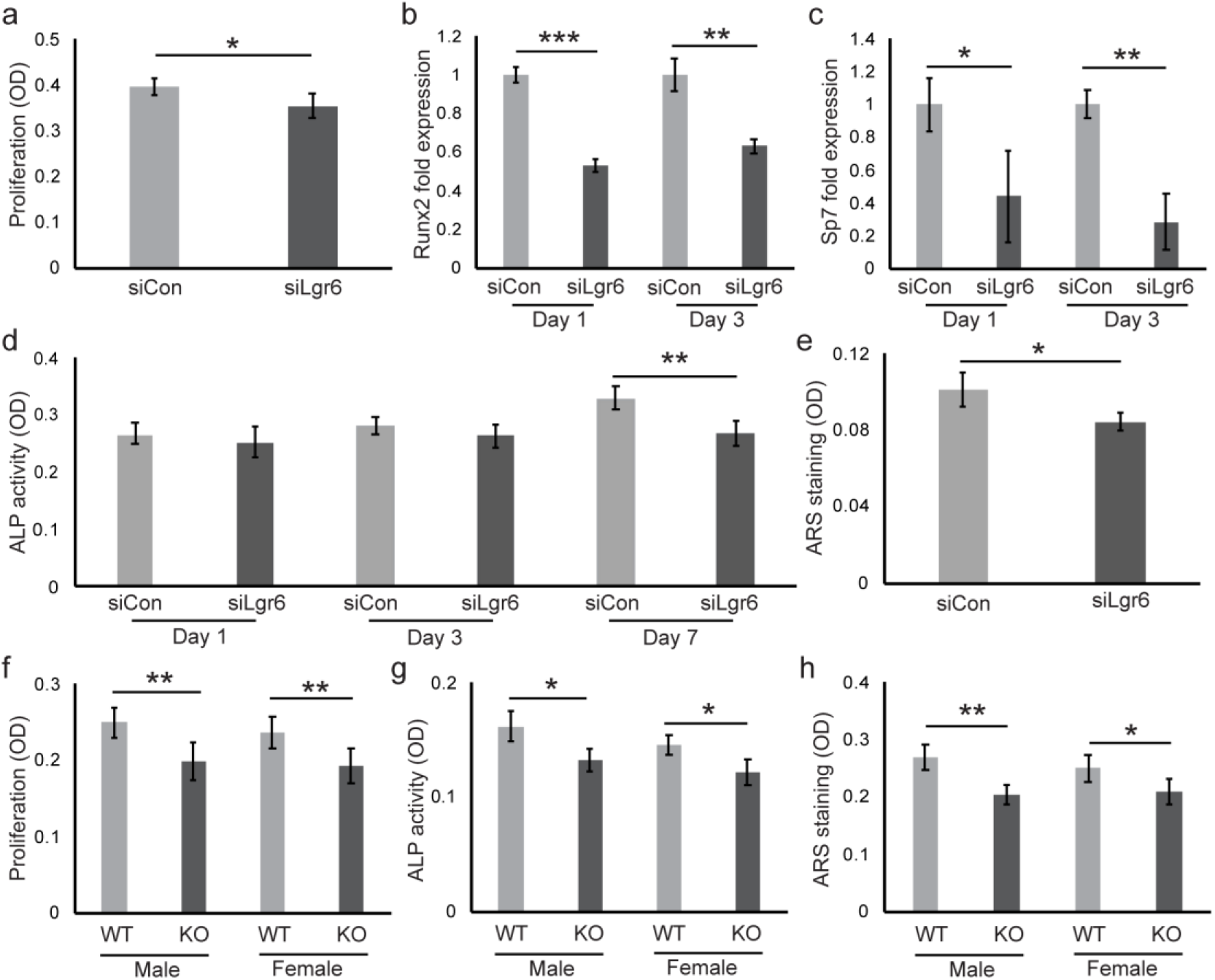
Knockdown of Lgr6 results in reduced osteoblast proliferation, differentiation, and mineralization in MSC derived osteoblasts. (a) Cell proliferation quantified by WST-1 assay in Lgr6 knockdown (siLgr6) and control knockdown (siCon) MSCs. (b and c) qPCR-derived RNA expression of Runx2 and Sp7 in Lgr6 knockdown MSCs relative to controls following 24 and 72 hours of osteoinduction. (d) Alkaline phosphatase (ALP) activity in knockdown MSCs 1, 3, and 7 days post-osteoinduction. (e) Quantification of alizarin red-S (ARS) staining of knockdown MSCs 18 days post-osteoinduction. Lgr6-KO and Lgr6-WT primary MSCs show reduced (f) proliferation by WST-1 assay, (g) ALP activity 12 days post-osteoinduction, and (h) ARS staining 21 days post-osteoinduction. Each experiment was performed in triplicate and analyzed by Student’s t-test; *p < 0.05, **p < 0.01, ***p < 0.001, ns = not significant. Error is reported as standard deviation.

To complement our Lgr6 transient knockdown experiments, we sought to evaluate the necessity of Lgr6 for MSC proliferation and osteodifferentiation using a genetic loss of function model. Using our newly generated Lgr6 mouse knockout allele (see Methods; Supplemental Figure 3), we generated primary MSCs from the femoral bone marrow of Lgr6 knockout (Lgr6-KO) and wildtype littermates (Lgr6-WT) and assessed their relative proliferation and osteogenic potential. Lgr6-KO cells had reduced proliferation as compared to Lgr6-WT MSCs (male=21%, p=0.002; female=18%, p=0.005) (Figure 1f). Using ex vivo osteodifferention assays to assess differentiation (day 11) and mineralization (days 18-21), we found Lgr6-KO cells had reduced alkaline phosphatase activity (male=19%, p=0.011; female=17%, p=0.013) and mineralization as compared to Lgr6-WT MSCs (male=23%, p=0.004; female=17%, p=0.045) (Figure 1g-h). The finding that knockdown of Lgr6 led to reduced osteodifferentiation led us to ask whether there was a broader change in MSC multipotency. Using an immortalized mouse MSC cell line, we evaluated the ability of C3H10T1/2 cells to undergo chondrogenesis or adipogenesis following Lgr6 siRNA-mediated knockdown. Quantification of oil red O staining 12 days post induction of adipogenesis and alcian blue staining 9 days post induction of chondrogenesis, revealed no significant difference between the Lgr6 knockdown and control cells (Supplemental Figure 4). These data suggest that Lgr6 is necessary for normal osteogenesis of MSCs, but not adipogenesis or chondrogenesis.

### Lgr6 overexpression enhances osteoblastogenesis in vitro

Having found a necessity for Lgr6 in osteoblastogenesis, we wanted to determine if overexpression of Lgr6 was sufficient to increase MSC proliferation or osteodifferentiation. We cloned an Lgr6 overexpression plasmid using the full-length mouse Lgr6 cDNA (pCAG-Lgr6, from here on referred to pLgr6). Following validation and transfection optimization (Supplemental Figure 5), we transfected pLgr6 and pCAG-GFP (control, from here on referred to pCAG) into MSCs and measured proliferation 24 hours post-transfection. Overexpression of Lgr6 resulted in a 20% increase in proliferation as compared to the control cells, as determined by WST-1 assay (p=0.033) (Figure 2a). To determine if any changes in osteodifferentiation were found with overexpression of Lgr6, osteogenic markers were assessed 7 days post-osteoinduction). While Lgr6 overexpression did not result in increased expression of early osteogenic markers Runx2 and Sp7 at this stage, it did result in significantly increased mRNA expression of later markers Alkaline phosphatase (75%, p=0.007) and Osteopontin (111%, p=3.89E-05) as compared to the pCAG controls (Figure 2b). Consistent with these data, Lgr6 overexpression led to increased ALP activity by day 7 (8%, p=0.012) (Figure 2c) and mineralization was increased at day 21 (10%, p=0.034) (Figure 2d). These findings were corroborated in experiments with primary mouse MCOs, whereby overexpression of Lgr6 led to significantly increased proliferation (14%, p=0.025) and increased ALP activity (8%, p=0.026) and mineralization (10%, p=0.025) 7 and 21 days post-osteoinduction, respectively (Supplemental Figure 6).

**Figure 2.**
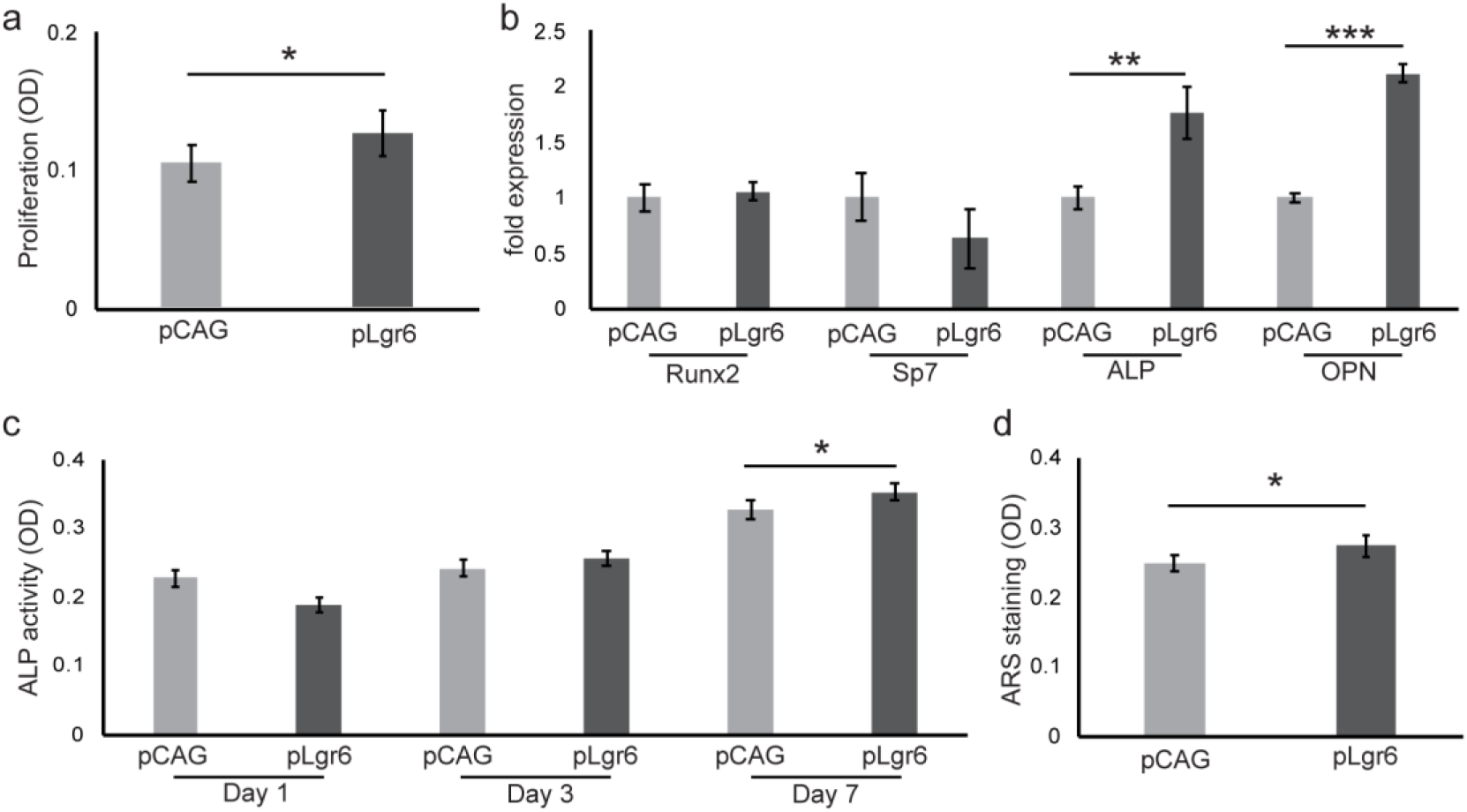
Lgr6 overexpression promotes MSC proliferation and osteodifferentiation. (a) Cell proliferation quantified by WST-1 assay in Lgr6 overexpression (pLgr6) and control (pCAG) MSCs. (b) qPCR-derived RNA expression of Runx2, Sp7, Alkaline phosphatase (ALP), and Osteopontin (OPN) in Lgr6 overexpression MSCs relative to controls following 7 days of osteoinduction. (c) ALP activity in Lgr6 overexpression MSCs 1, 3, and 7 days post-osteoinduction. (d) Quantification of ARS staining of Lgr6 overexpression MSCs 18 days post-osteoinduction. Each experiment was performed in triplicate and analyzed by Student’s t-test; *p < 0.05, **p < 0.01. Error is reported as standard deviation.

### Lgr6 is necessary to attain peak bone mass in mice

To gain in vivo context to our MSC derived data, we evaluated mouse long bones for the cellular localization of LGR6 expression. Using the Lgr6 GFP reporter mouse allele^12^ (Lgr6-EGFPcreERT), we evaluated 2-month-old femoral trabeculae by anti-GFP immunohistochemistry. This staining reveals Lgr6-GFP cells at the periphery of the trabecular bone marrow (Figure 3a and b). The location and sporadic expression pattern of Lgr6-GFP cells could be consistent with either osteoblast or osteoclast progenitors, however we do not find them to co-label with TRAP, indicating they are not osteoclasts (Figure 3a-a’’). Conversely, anti-SP7 staining reveals a subset of SP7-expressing cells are also Lgr6-GFP positive, supporting the data that Lgr6 is a marker of osteoprogenitors (Figure 3b>-b’’). To determine if these Lgr6-expressing cells give rise to additional cells in the bone during homeostasis, we performed a genetic lineage analysis using the Lgr6 tamoxifen inducible cre allele (Lgr6-EGFPcreERT) bred to a tdTomato cre reporter allele (Ai9)^12,28^. 2-month-old compound heterozygous mice were tamoxifen induced to genetically mark Lgr6-expressing cells with tdTomato. Five days post-tamoxifen, Lgr6-expressing cell descendants are present on the surface of the femoral trabeculae of bone. (Figure 3c). 17 days post-tamoxifen, Lgr6-expressing cell descendants can occasionally be found in the adjacent cortical bone (Figure 3d). Taken together, these expression and lineage studies support Lgr6-expressing cells in the bone marrow as osteoprogenitors that contribute to bone homeostasis.

**Figure 3.**
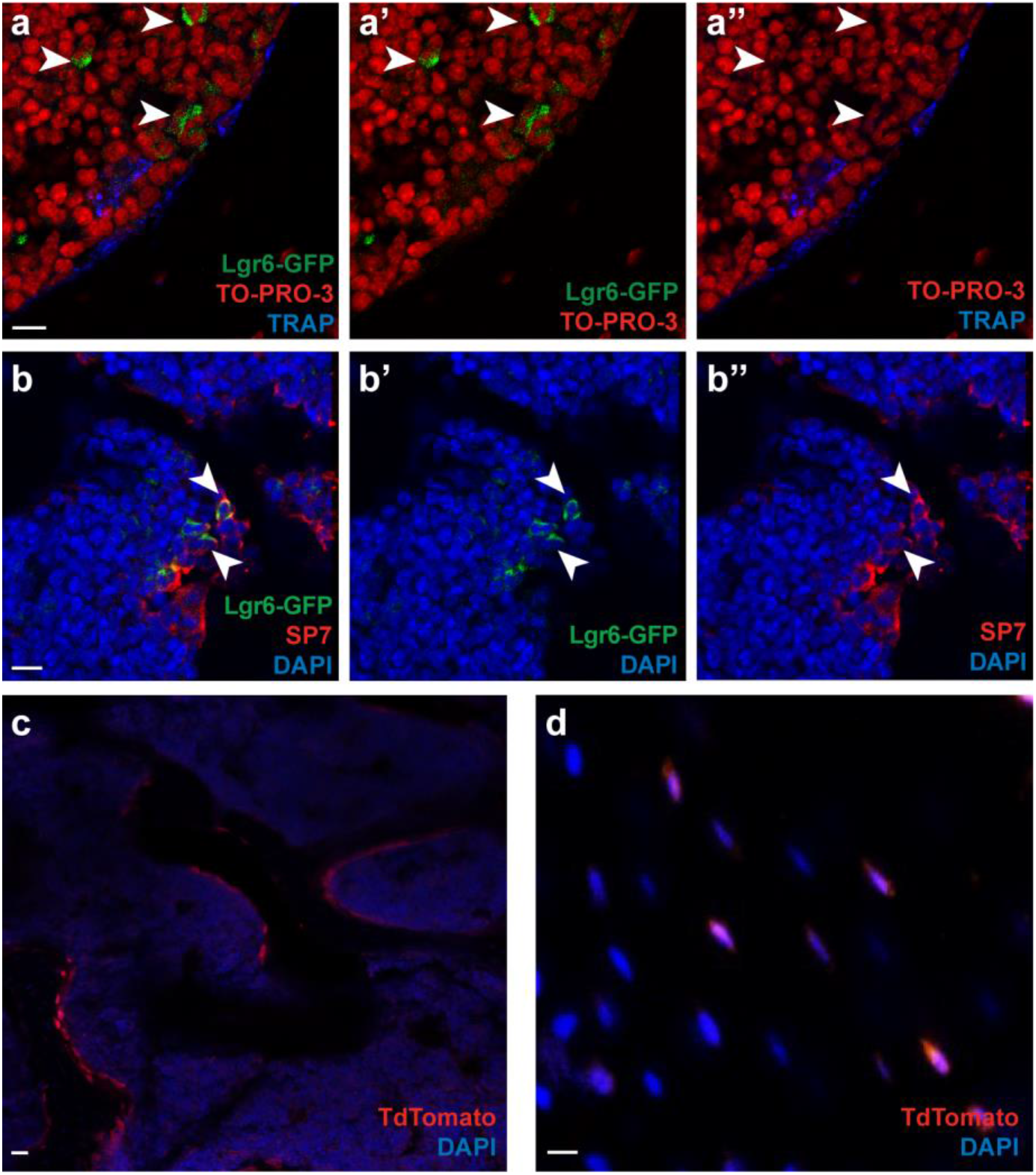
Immunohistochemistry and genetic lineage analysis of Lgr6-expressing osteoprogenitors. (a and b) Immunohistochemistry of adult Lgr6-EGFPcreERT2 mouse femur with sections of trabecular bone from femoral head. (a-a”) Few Lgr6 expressing cells (green; marked by arrowheads) are found within the bone marrow and do not co-label with TRAP stained osteoclasts (blue). Nuclei are stained with To-Pro-3 (red). (b-b”) Lgr6-expressing cells (green; marked by arrowheads), co-label with subset of Sp7 expressing osteoblasts (red) found at the periphery of the trabeculae. Nuclei are stained with DAPI (blue). (c and d) Immunohistochemistry of adult tamoxifen induced Lgr6-EGFPcreERT2;R26R-CAG-LSL-TdTomato mouse femur with section through (c) femoral head trabeculae and (d) cortical bone; nuclei are marked with DAPI (blue). (c) Lgr6-expressing cell descendants (red) are found surrounding the trabeculae 5 days post-tamoxifen induction, and (d) on the endosteal surface of the adjacent cortical bone 17 days post-tamoxifen induction. Scale bars are 10µm.

We next sought to determine whether Lgr6 is necessary for bone homeostasis. We analyzed 8-week-old male and female femurs from Lgr6-KO and Lgr6-WT mice by micro computed tomography (μCT) (Figure 4). Analysis of the bone parameters revealed that the distal femurs of Lgr6-KO mice had decreased trabecular bone volume as compared to Lgr6-WT mice (BV/TV, male=22%, p=0.013; female=22%, p=0.046) (Figure 4b). Analysis of additional trabecular parameters reveal a 7% decrease in trabecular thickness in female Lgr6-KO mice (Tb.Th, p=0.037), as well as a 14% increase in trabecular separation (Tb.Sp, p=0.019) and a 13% decrease in trabecular number (Tb.N, p=0.006) in male Lgr6-KO mice (Figure 4c-e). μCT analysis of the mid-cortical segment of Lgr6-KO and Lgr6-WT femurs did not find any significant differences (Supplemental Figure 7).

**Figure 4.**
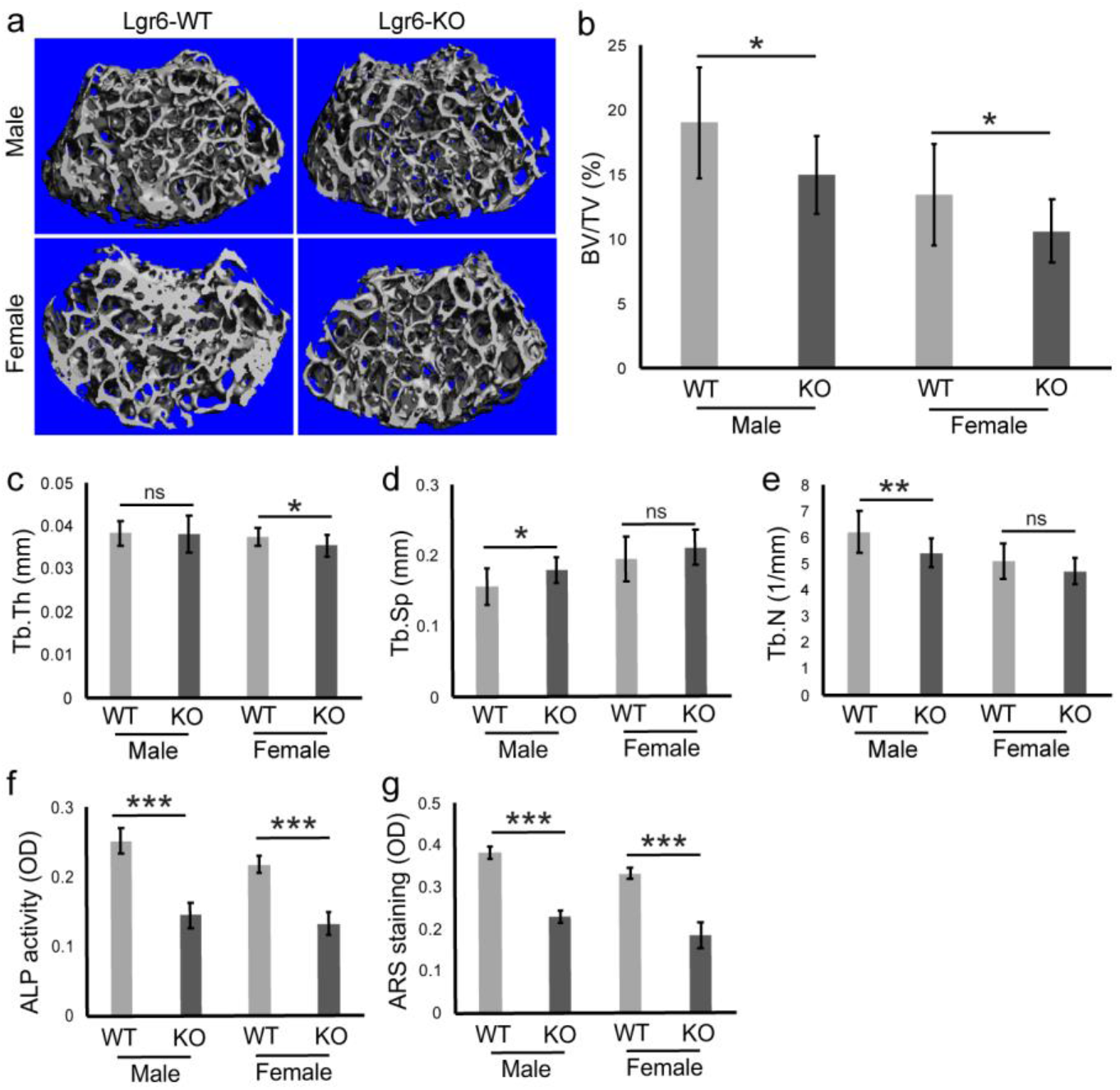
Lgr6 knockout mice have decreased trabecular bone parameters compared to controls. MicroCT analysis of femurs from 8-week-old Lgr6 wildtype (WT) and Lgr6-knockout (KO) male and female mice. (a) Representative 3D images of trabeculae from each sex and genotype. (b-e) Trabeculae bone parameter with significant differences between genotypes: (b) bone volume/tissue volume (BV/TV), (c) trabecular thickness (Tb.Th), (d) trabecular separation (Tb.Sp), and (e) trabecular number (Tb.N). Cohort sizes: 14 female Lgr6-WT mice, 11 female Lgr6-KO mice, 11 male Lgr6-WT mice, and 12 male Lgr6-KO mice. Quantitative bone parameters were analyzed for significance by Student’s t-test; sexes were analyzed separately. (f and g) Osteodifferentiation of primary bone marrow cells from Lgr6-WT and Lgr6-KO littermates. (f) Measurement of ALP activity 12 days of post-osteoinduction. (g) Quantification of alizarin red-S (ARS) staining 21 days post-osteoinduction. Each bone marrow cell osteodifferentiation experiment was performed in triplicate and analyzed by Student’s t-test; *p < 0.05, **p < 0.01, ***p < 0.001, ns = not significant. Error is reported as standard deviation.

To further explore the function of Lgr6 in osteogenesis, we used ex vivo bone marrow culture from Lgr6-KO and Lgr6-WT mice and induced osteogenesis. 11 days post-osteoinduction, Lgr6-KO cells showed significantly reduced ALP activity (male=30%, p=0.0001; female=22%, p=0.0001) (Figure 4f). 18-21 days post-osteoinduction, Lgr6-KO cells had reduced mineralization as compared to Lgr6-WT cells of the same sex (male=32%, p=5.203E-06; female=37%, p=0.0001) (Figure 4g). Taken together, these data suggest that the Lgr6-KO trabecular bone phenotype is due to the reduced differentiation capability of Lgr6-expressing progenitor cells towards osteoblasts.

### RSPO2 and MaR1 differentially regulate LGR6 receptor signaling in osteoblasts

Having defined a role for Lgr6 in osteoblast differentiation and bone homeostasis, we next addressed LGR6 receptor signaling in osteoprogenitors. LGR6 has two defined ligands, R-spondins (RSPO1-4) which when bound agonize WNT signaling, and maresin-1 (MaR1) which potentiates GPCR signaling as has been described in phagocytes^19,20^ ; these ligand-receptor-signaling interactions have not yet been defined in osteogenic cells. Of the four R-spondins, RSPO2 binds to LGR6 with the highest affinity in vitro and has an established role in osteoblastogenesis and bone development, thus RSPO2 is the R-spondin ligand we focused on in our experiments^22,23^. We first assessed WNT/β-catenin signaling using the TOPFlash luciferase assay. Using C3H10T1/2 cells with their endogenous level of Lgr6, MaR1 did not significantly stimulate WNT signaling, but RSPO2 did, as is found in other cell types^19,29^ (102% increase, p=0.005) (Figure 5a). These findings were corroborated in the context of Lgr6 overexpression. Overexpression of Lgr6 in C3H10T1/2 cells resulted in increased WNT signaling (pBKS vs. pLgr6, 46%, p=9.8E-4), which was further agonized by RSPO2 (pLgr6 vs. pLgr6/RSPO2, 34%, p=7.0E-4) (Figure 5b). While no significant difference was found between pLgr6 plus RSPO2 versus the RSPO2 control (pBKS/RSPO2 vs. pLgr6/RSPO2, p=0.0515), this experiment was confounded by endogenous expression of Lgr6 in C3H10T1/2 cells, as well as the expression of Lgr4 which also agonizes WNT signaling upon RSPO binding in vitro^30,31^. Intriguingly, overexpression of Lgr6 in the presence of MaR1 inhibits pLgr6-induced WNT signaling (pLgr6 vs. pLgr6/MaR1, p=3.8E-4) suggesting reciprocal roles for MaR1 and RSPO2 in LGR6 mediated WNT signaling (Figure 5b).

**Figure 5.**
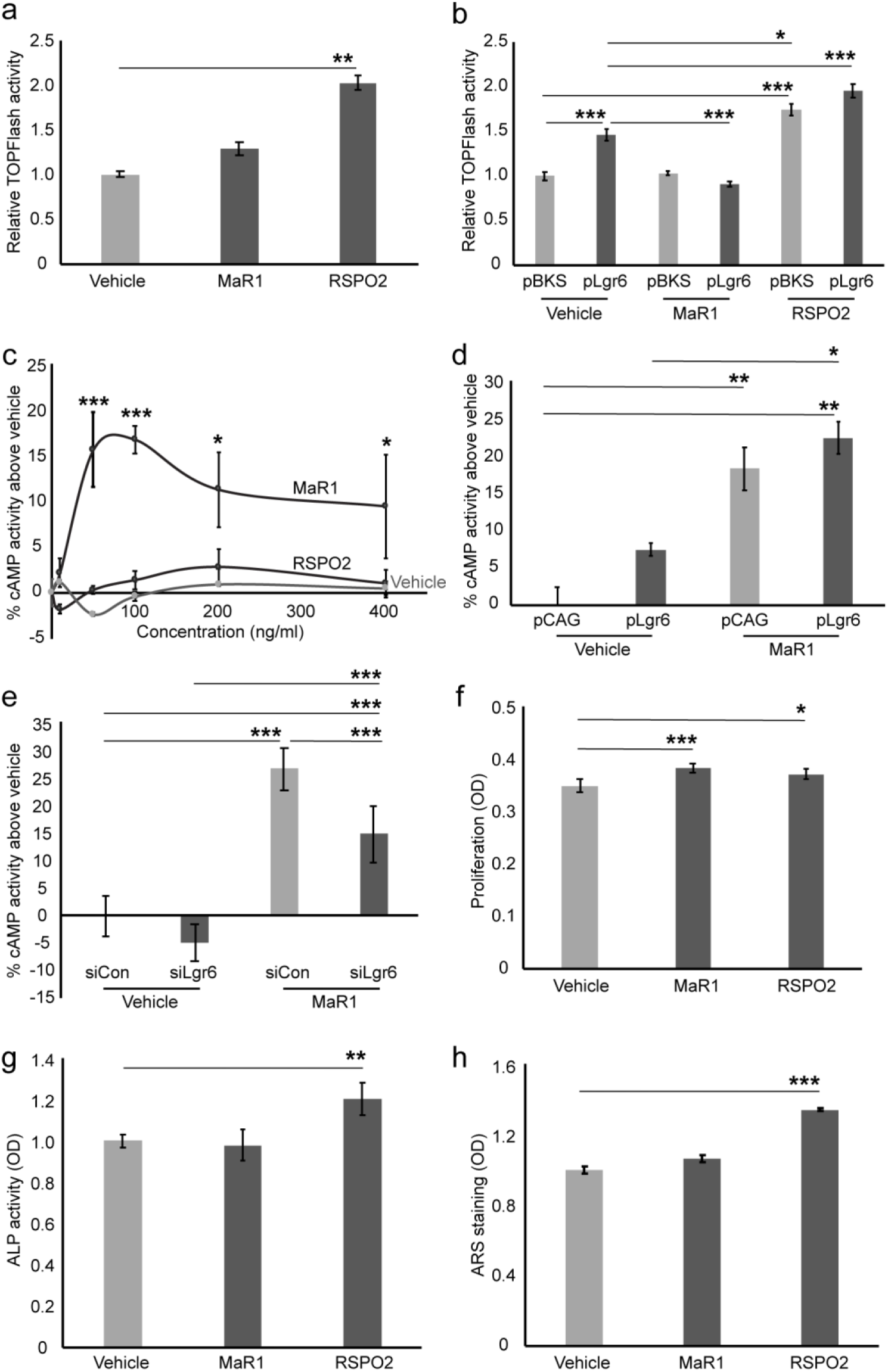
Lgr6 mediated WNT/β-catenin and GPCR/cAMP signaling in response to RSPO2 or MaR1 in C3H10T1/2 cells. (a and b) TOPFlash luciferase assay to assess WNT/β-catenin signaling. (a) WNT signaling in the presence of 100ng/ml MaR1 or 100ng/ml RSPO2 as compared to vehicle control. (b) WNT signaling in the context of Lgr6 overexpression (pLgr6) or control (pBKS) when exposed to 100ng/ml MaR1, 100ng/ml RSPO2, or vehicle control. (c-e) cAMP activity assay to assess GPCR/cAMP signaling. (d) cAMP activity at over a range of MaR1 and RSPO2 concentrations (0-400ng/ml). (d) cAMP activity in the context of Lgr6 overexpression (pLgr6) or control (pCAG) when exposed to 100ng/ml MaR1 or vehicle control. (e) cAMP activity in the context of Lgr6 knockdown (siLgr6) or control (siCon) when exposed to 100ng/ml MaR1 or vehicle control. All signaling experiments were performed in technical duplicate and biological triplicate, and analyzed by one-way ANOVA test with Tukey’s HSD posthoc analysis. (f-h) Proliferation and osteodifferentiation assays using in the presence of 100ng/mL MaR1, 100ng/ml RSPO2, or vehicle control. (f) Cell proliferation measured by WST-1 assay, (g) ALP activity 3 days post-osteoinduction, and (h) ARS staining 21 days post-osteoinduction. Each experiment was performed in triplicate and analyzed by Student’s t-test, *p < 0.05, **p < 0.01, ***p < 0.001. Error is reported as standard deviation.

We next evaluated the ability of RSPO2 and MaR1 to stimulate G-protein signaling by assaying cAMP activity in C3H10T1/2 cells. While no concentration of RSPO2 induced cAMP activity above the level of the vehicle control, MaR1 treatment led to an increase in cAMP activity at all concentrations tested greater than or equal to 50 ng/ml (50 ng/ml: p=1.2E-5, 100 ng/ml: p=2.5E-5, 200 ng/ml: p=0.01, 400 ng/ml: p=0.035) (Figure 5c). To determine whether MaR1-induced cAMP activity is Lgr6 specific, we evaluated this signaling in the context of Lgr6 knockdown and overexpression. Consistent with the dose curve data, treatment of cells with 100 ng/ml MaR1 in the context of control or Lgr6 overexpression resulted in increased cAMP activity (pCAG/vehicle vs. pCAG/MaR1, p=0.0046; pCAG/vehicle vs. pLgr6/MaR1, p=.0022). While no significant difference in cAMP activity was found between MaR1-treated pCAG and pLgr6 expressing cells, there is an increase in signaling in Lgr6 expressing cells between vehicle and MaR1 treatment (pLgr6/vehicle vs. pLgr6/MaR1, p=0.011), suggesting Lgr6 dependency (Figure 5d). In complementary experiments, siRNA-mediated knockdown of Lgr6 in the presence of MaR1 results in a 55% reduction in cAMP activity as compared to MaR1 treated control siRNA cells (p=3.1E-4) (Figure 5e). These results demonstrate that MaR1-induced cAMP activity in C3H10T1/2 cells is, at least in part, specific to Lgr6.

To determine whether exogenous RSPO2 and MaR1 influence osteogenesis, we performed proliferation and osteodifferentiation assays with C3H10T1/2 cells. Analysis of proliferation by WST-1 assay determined that both MaR1- and RSPO2-treated cells had increased proliferation (18%, p=0.0008 and 33%, p=0.02, respectively) (Figure 5f). Following induction of osteodifferentiation, RSPO2-treated cells had increased ALP activity 7 days post-induction (17%, p=0.006) and increased mineralization 21 days post-induction (7%, p=0.0002) as compared to vehicle controls (Figure 5g-h). Interestingly, while MaR1 treatment did increase proliferation, it did not influence osteodifferentiation (Figure 5g-i), suggesting MaR1 and RSPO2 differentially regulate Lgr6 during osteogenesis.

## DISCUSSION

Lgr6 is a marker of osteoprogenitor cells and is dynamically expressed during osteoblastogenesis such that it is highly expressed (Lgr6^high^) as cells are proliferating and more lowly expressed (Lgr6^low^) as cells differentiate^14,16,17^. In our present study, we used in vitro loss of function experiments in mouse primary cells (MSCs and MCOs) and demonstrated that loss of Lgr6 results in decreased proliferation, differentiation, and mineralization, which we also found with Lgr6-KO MSCs. In complementary experiments we found that overexpression of Lgr6 leads to increased proliferation, differentiation, and mineralization of osteoprogenitors. While these data clearly demonstrate a role for Lgr6 in osteoblastogenesis, it raises the question as to when LGR6 exerts its function during the process. Does Lgr6 regulate osteoprogenitor proliferation when its expression is high (Lgr6^high^) with an indirect effect on osteodifferentiation or does Lgr6 directly regulate both processes during osteoblastogenesis (ie. Lgr6^high^ regulates proliferation, Lgr6^low^ regulates differentiation)? Our signaling data support the latter model. We have demonstrated for the first time in osteoprogenitors, that RSPO2/LGR6 enhances WNT signaling and MaR1/LGR6 stimulates cAMP activity, indicating that LGR6 can mediate two different signaling pathways in a single cell type. Moreover, we find that RSPO2 increases osteoprogenitor proliferation and differentiation, whereas MaR1 only significantly increases proliferation, not differentiation, further supporting our model. Within the framework of this two ligands/one receptor model, it remains to be determined how in a single cell type, RSPO2 and MaR1 regulate two different pathways via LGR6 receptor binding. The finding that MaR1 inhibits Lgr6 induced WNT signaling (Figure 5b, Vehicle/pLgr6 vs. MaR1/pLgr6) suggests competitive ligand binding, which has yet to be explored. However, the true signaling model could be more complex. Our data reveal modest RSPO2-induced WNT signaling and MaR1-induced cAMP activity even without over-expression of Lgr6 (Figures 5b and 5d, respectively). This finding may simply be due to endogenous background expression of LGR6 in C3H10T1/2 cells, or it could reflect LGR6-dependent and -independent roles for both ligands. It is well-established that RSPO1, RSPO2, RSPO3, and RSPO4 are all capable of binding receptors LGR4, LGR5, and LGR6 to agonize WNT signaling^19,30^. Because LGR4 is also expressed in osteoprogenitors^14,32^, RSPO2 (and potentially the other R-spondins) may redundantly agonize WNT signaling through LGR4. The G-protein signaling role for MaR1 beyond its interactions with LGR6 is less clear. Orphan G-protein coupled receptor screening with MaR1 identified specific signaling with LGR6, not LGR4 or LGR5^33^, indicating that any LGR6-independent MaR1 induced cAMP activity (Figure 5d, MaR1/pCAG and Figure 5e, MaR1/siCon) would likely be indirect or utilizing an as-of-yet unidentified receptor.

Importantly, our in vitro Lgr6 data are supported by our in vivo mouse experiments. By histology we demonstrate that Lgr6-expressing cells sparsely populate the femoral trabecular bone marrow and genetic lineage analyses reveal that Lgr6-expressing cell descendants contribute to the cells of the trabecular surface and express SP7, supporting that Lgr6-expressing cells are osteoblastic. Consistent with these findings, genetic loss-of-function experiments reveal significantly reduced BV/TV and trabecular thickness and increased trabecular separation in Lgr6-KO mice compared to Lgr6-WT controls. It remains to be determined whether increased LGR6 activity in MSCs in vivo would result in increased bone production like we find in vitro (Figure 2 and Supplemental Figure 6), which could provide inroads to novel anabolic therapies. While the result of exogenous RSPO2 treatment on bone has not been reported, exogenous MaR1 has been associated with bone regeneration^24,34^. Exogenous MaR1 can stimulate bone formation in rat molar sockets and resolve macrophage based inflammation resulting in improved fracture healing in aged mice, though these studies did not explore LGR6 signaling^24,34^. Moving forward it will be important to determine if RSPO2 and/or MaR1 treated mice have increased bone mass, and whether this is an Lgr6-specific phenotype. Furthermore, our focus on the role of Lgr6 in bone originated in our finding that Lgr6 is expressed in the mouse digit tip bone and is necessary for digit tip bone regeneration following amputation^13^. While this present study focuses on a broader role for Lgr6 in bone homeostasis, our prior digit tip studies as well as the MaR1 bone regeneration studies suggest that Lgr6 also has an important role in bone fracture repair which would be an important focus of future studies^13,24,34^.

## MATERIALS AND METHODS

### Mice

All mouse breeding and experimentation was done with the approval of the Brigham and Women’s Hospital IACUC. Primary MSCs and MCOs were derived from wildtype C57BL/6 mice (JAX #000664; The Jackson Laboratory (JAX), Bar Harbor, ME, USA) from a specific pathogen free in-house breeding colony. Lgr6 expression studies used 8-week-old female Lgr6-EGFPcreERT2 mice (JAX #016934). Lgr6 genetic lineage analyses used the Lgr6-EGFPcreERT2 inducible cre allele and the R26R-CAG-LSL-tdTomato (Ai9; JAX #007909) cre reporter allele^12,28^. 8-week-old compound heterozygous Lgr6-EGFPcreERT2;R26R-CAG-LSL-tdTomato heterozygous mice were given three consecutive daily doses of 3mg/40g body weight tamoxifen in corn oil by oral gavage. Mice were euthanized and bones were collected for immunohistochemistry on day 5 or day 17 post-tamoxifen induction.

The Lgr6 knockout allele (Lgr6-KO) was generated secondary to our efforts to engineer a conditional floxed allele. Using the dual guide CRISPR-Cas9 genome modification strategy^35–37^, sgRNAs were designed to Lgr6 intron 14 and intron 16 on mouse chromosome 1 (Supplemental Figure 3, Supplemental Table 1); deletion of exons 15 and 16 results in frameshift and a nonsense transcript. Editing efficiency was validated in NIH3T3 cells, then crRNAs were commercially synthesized, as were two loxP-containing targeting oligonucleotides (IDT, Coralville, IA, USA). Guide sequences were Lgr6_16.2 AltR1-ACAGAUCAUACAUUUCUCAGGUUUUAGAGCUAUGCU-AltR2 and Lgr6_14.3 AltR1-GGCCAAGACCGGGAGCCUUGGUUUUAGAGCUAUGCU-AltR2; tracrRNA was ready made from IDT. The Harvard Genome Modification Facility generated RNPs with the crRNA:tracrRNA guides and Cas9 nuclease (IDT) and microinjected them with the targeting oligonucleotides into fertilized C57BL/6J mouse eggs. (Oligos were included in the injection in attempts to generate a floxed allele; the null allele described here does not have integration of these oligos.) Genomic DNA from all progeny was screened by PCR amplicon sequencing for integration of loxP sequences, and by long range PCR product cloning and sequencing to capture any locus deletions. For our new knockout allele (Lgr6-KO), a single founder from the above Lgr6 CRISPR-Cas9 microinjection was found to have a deletion of exons 15 and 16 (without introduction of loxP sequences). PCR amplicon DNA sequencing found no mutations in putative exonic off-target cut sites on mmChr1; genomic and qPCR analyses verify this to be a null allele (Supplemental Figure 3). The founder mouse was bred one generation onto FVB/NJ wildtype background, then crossed back into the C57BL/6 wildtype background for four generations. The Lgr6-KO mice are homozygous viable and fertile, with no overt phenotypes, as is found for the published Lgr6-null allele^12^.

### Cell culture

8-week-old female wildtype mice were used for isolation of primary MSCs by standard protocols^38–40^. In brief, femurs were dissected and cleaned of soft tissue in PBS. Femoral ends were removed, and bone marrow cells were flushed out with a 22G needle syringe and collected and plated for tissue culture in α-MEM supplemented with 10% FBS, 2mM L-glutamine, and 1% penicillin/streptomycin. Adherent cells were grown to confluency and were repeatedly trypinized and passaged eight times. The resultant cells were used as MSCs in experiments as described. For ex vivo bone marrow osteoblast culture, bone marrow was isolated from 8-week-old female Lgr6-WT or Lgr6-KO mice as described above and used directly in osteodifferentiation assays without generation of adherent MSCs.

Primary mouse calvarial osteoblast cells (MCOs) were isolated by standard protocols^14,39,40^. Postnatal day one wildtype mouse pups were euthanized, calvaria were dissected from skulls, cleaned of soft tissue, cut into small pieces, and subjected to five sequential dispase/collagenase digestions. Cells liberated in the second to fifth digestions were pooled and placed in FBS to neutralize the dissociation enzymes. Cells were concentrated and plated for tissue culture in proliferative media (α-MEM supplemented with 10% FBS, 2mM L-glutamine, and 1% penicillin/streptomycin) and used as MCOs in experiments as described.

C3H10T1/2 cells (ATCC CCL-226, American Type Culture Collection, Manassas, VA, USA), immortalized mouse MSCs, were obtained from the Bone Cell Core at the MGH Center for Skeletal Research. C3H10T1/2 cells were in maintained in proliferative tissue culture medium (α-MEM supplemented with 10% FBS, 2mM L-glutamine, and 1% penicillin/streptomycin) and used for experiments as described.

### In vitro siRNA-mediated knockdown

The TriFECTa DsiRNA system (IDT) was used for siRNA experiments; predesigned siRNAs were used for Lgr6 (siLgr6: mm.Ri.Lgr6.13) and negative control (siCon: DS NC1). For Lgr6 knockdown experiments, 1×10^4^ primary MCOs or MSCs were seeded in 24-well plates and cultured for 24 hours in proliferative media. At 40-50% confluency, cells were transfected with siLgr6 or siCon siRNAs (100nM or as noted) using the Lipofectamine RNAiMAX transfection reagent (Thermo Fisher, Waltham, MA, USA) according to the manufacturer’s protocol. Subsequent cell culturing and treatment is as noted for individual assays. Lgr6 protein levels were analyzed by standard western blot analysis using anti-LGR6 (ab126747, 1:1000; Abcam, Cambridge, MA, USA) and anti-GAPDH (ADI-CSA-335-E, 1:5000; Enzo Life Sciences, Farmingdale, NY, USA) antibodies. All knockdown experiments were performed in technical duplicate and biological triplicate.

### in vitro cDNA overexpression

The Lgr6 overexpression plasmid (pCAG-Lgr6) was constructed by replacing the GFP cassette in the control vector (pCAG-GFP; plasmid #11150, Addgene, Cambridge, MA, USA) with the mouse Lgr6 coding sequence. Bases 137 to 3040 from NCBI genbank accession NM_001033409.3 was amplified from mouse MSC-derived cDNA, PCR purified, and cloned into the NotI linearized pCAG backbone using Gibson assembly (NEB, Ipswich, MA, USA).

The resultant plasmid was DNA sequenced and no mutations were found. For overexpression experiments, 1×10^4^ MCOs, MSCs, or C3H10T1/2 cells were plated in 24-well plate and cultured for 24h in proliferative media. At 40-50% confluency, cells were transfected pCAG-Lgr6 or control plasmid (100 ng/ml or as noted) using the Fugene HD transfection reagent (Promega, Madison, WI, USA). Subsequent cell culturing and treatment is as noted for individual assays. All overexpression experiments were performed in technical duplicate and biological triplicate.

### Quantitative PCR

Following MSC or MCO Lgr6 knockdown or over-expression experiments, cells to be analyzed by qPCR were collected and total RNA was extracted using the Qiagen RNeasy Plus Micro kit (Qiagen, Germantown, MD, USA). First strand cDNA was synthesized using SuperScript IV reverse transcriptase with oligo dT primers (Thermo Fisher). Purified cDNA from experimental samples was used as input for qPCR. Using the QuantStudio 5 Real-Time PCR 384-well system (Applied Biosystems, Foster City, CA, USA), osteogenic (Sp7, Runx2, ALP, and OPN) and control (HPRT) transcripts were amplified (Supplemental Table 1) using SsoAdvanced Universal SYBR Green Supermix (Biorad, Hercules, CA, USA). Established TaqMan assays were used to determine the expression of Lgr6 and TBP intra-well controls. Delta threshold cycle (ΔCt) was calculated for each well using the 2^−ΔCt^ method^41^. All qPCR experiments were performed in both technical and biological triplicate.

### Cell proliferation assay

The WST-1 reagent (Sigma-Aldrich, St. Louis, MO, USA) was used for all experiments assessing cellular proliferation. MCOs, MSCs, or C3H10T1/2 transfected cells (as detailed above), or Lgr6-KO and control cells were cultured for 24 hours in proliferative media. 10uL of WST-1 reagent was added to the wells four hours prior to harvest. Following the incubation, all wells were analyzed on a colorimetric plate reader at 450nm (BioRad). All proliferation assays were performed in technical duplicate and biological triplicate.

### In vitro differentiation assays

For osteodifferentiation assays, cells growing in proliferative media were changed into osteogenic media which was changed every 48 hours. For C3H10T1/2 cells and primary MSCs, osteogenic media composition was α-MEM supplemented with 10% FBS, 2mM L-glutamine, 10mM β-glycerophosphate, 50 μg/mL ascorbic acid, 100nM dexamethasone, and 1% penicillin/streptomycin; for primary MCOs, osteogenic media composition was α-MEM supplemented with 10% FBS, 2mM L-glutamine, 10mM β-glycerophosphate, 50 μg/mL ascorbic acid and 1% penicillin/streptomycin^14,38–40^. Length of osteodifferentiation and choice of terminal assay varied by experiment as detailed throughout. For experiments assaying alkaline phosphatase (ALP) activity, cells were in PBS, then subjected to a -80°C/37°C freeze-thaw cycle. Para-nitrophenylphosphate (PNPP) substrate was used to measure ALP activity by colorimetric quantification at 405nm. For experiments assaying alizarin red-S staining, cells were harvested 18-21 days post-osteoinduction. Cells were fixed in 4% paraformaldehyde and stained with 40mM Alizarin red-S. For quantification, cells were incubated in 10% acetic acid and collected by scraping. Samples were incubated at 90°C for 5 min followed by centrifugation; resultant supernatant was mixed with 10% ammonium hydroxide and used for colorimetric quantification at 405 nm^42^. For adipogenic differentiation, cells are incubated for 12 days in adipogenic induction medium containing 100nM dexamethasone, 0.5µM isobutylmethylxanthine (IBMX) and 10ng/ml insulin, with media changes every 48 hours. Oil Red O was used to stain adipocytes and was quantified at 510nm following isopropanol dye extraction^43^. For chondrogenic differentiation, cells were cultured for 12 days in high-glucose DMEM supplemented with 100nM dexamethasone, 1% insulin-transferrin-sodium selenite (ITS), 50µM ascorbate-2-phosphate, 1mM sodium pyruvate, 50µg/ml proline, and 20ng/ml TGFβ3 for 12 days. Alcian blue was used to stain chondrocytes and was quantified at 620nm following hydrochloric acid extraction^44^.

### Immunohistochemistry

Femurs from Lgr6-EGFPcreERT2 mice were collected in PBS, cleaned of soft tissue, and then fixed in 4% PFA overnight at 4°C. The bones were washed in PBS, taken through an increasing sucrose gradient (5-30%), then embedded and frozen in OCT freezing medium (Sakura Finetek, Torrance, CA, USA), and cut into 16-20µm cryosections. Sections were blocked in 5% goat serum then incubated with primary antibody at 4°C overnight (anti-dsRed for tdTomato (632496, 1:1000; Takara Bio, Mountain View, CA, USA), anti-SP7 (ab22552, 1:2000; Abcam), anti-GFP (ab13970, 1:1000; Abcam)). Sections were washed with PBST and incubated with species-specific Cy3, or A647 fluorophore-conjugated secondary antibody (Jackson Immunoresearch, West Grove, PA, USA) at room temperature for 1 hour. Slides were washed in PBST, counterstained with DAPI or TO-PRO-3 (Thermo Fisher), and coverslipped with Fluoromount-G mounting medium (Southern Biotech, Birmingham, AL, USA). TRAP staining was performed using the ELF-97 endogenous phosphatase detection kit (Thermo Scientific). A Zeiss LSM800 confocal microscope was used for fluorescent imaging at the BWH Confocal Microscopy Core.

### Micro computed tomography

Femurs were isolated from 8-week-old Lgr6-WT and Lgr6-KO mice and stored in 70% ethanol prior to analysis. For microCT analysis, a Scanco Medical μCT 35 system with an isotropic voxel size of 7μm was used to image femurs. Scans were conducted in 70% ethanol using an X-ray tube potential of 55 kVp, an X-ray intensity of 0.145 mA and an integration time of 400 ms. A region starting at a reference point anchored 0.35 mm proximal to the growth plate and extending 1.4 mm proximally was selected for distal femur trabecular bone analysis. A second region 0.56 mm in length located at the midshaft of the femur was used to calculate cortical parameters. A semi-automated contouring approach was used to distinguish cortical and trabecular bone. The region of interest was thresholded using a global threshold that set the bone/marrow cut-off at 453 mg HA/ccm for trabecular bone and 772 mg HA/ccm for cortical bone. 3-D microstructural properties of bone, including bone volume fraction (BV/TV), trabecular thickness (Tb.Th), trabecular number (Tb.N), trabecular separation (Tb.Sp) were calculated using software supplied by the manufacturer. Bone parameters are reported according to standard guidelines^45^.

### WNT signaling luciferase reporter assay

1×10^4^ MSCs or C3H10T1/2 cells were seeded in 24-well plates, grown to 40-50% confluency for transfection. All wells were transfected with the 100ng TOPFlash WNT signaling reporter plasmid^46^ (M50 Super 8x TOPFlash was a gift from Randall Moon; Addgene plasmid #12456) and 20ng pRL-TK Renilla control plasmid construct (Promega) using Fugene HD transfection reagent. Cells were co-transfected with 100ng pCAG-Lgr6 overexpression plasmid (pLgr6) or pBluescript II KS+ (pBKS) (Agilent Technologies, Santa Clara, CA, USA) as a control. RSPO2 (100ng/ml or as noted) (R&D Systems, Minneapolis, MN, USA), MaR1 (100ng/ml or as noted) (Cayman Chemical, Ann Arbor, MI, USA), or vehicle control (0.1% ethanol) was added to the media of the transfected cells. 72 hours post-transfection, cells were harvested for luciferase and renilla quantification using the Dual-Luciferase Reporter Assay System (Promega). Luciferase activity divided by renilla activity was used to calculate TOPFlash activity. Experiments were performed technical duplicate and biological triplicate.

### cAMP activity assay

1×10^4^ MSCs or C3H10T1/2 cells were seeded in 24-well plates, grown to 40-50% confluency for transfection. In overexpression experiments, cells were transfected with 100ng pCAG-Lgr6 (pLgr6) or pCAG-GFP (pCAG) control; in knockdown experiments, cells were transfected with siLgr6 or siCon control. 24 hours post-transfection, samples cells were treated with RSPO2 (100ng/ml or as noted), MaR1 (100ng/ml or as noted), or vehicle control (0.1% ethanol) in cAMP induction media for 30 minutes. Following the treatment, cells were harvested for quantification of cAMP activity using the cAMP-GLO Assay Kit (Promega). Percent increase in cAMP activity was calculated by the difference between control and treated groups, divided by control, then converted into a percentage. Experiments were performed in technical duplicate and biological triplicate.

## ACKNOWLEDGEMENTS

We thank the Confocal Microscopy Core at BWH for providing Zeiss LSM800 confocal access and the Harvard Genome Modification Facility for mouse allele generation. Support for this research was provided by a grant from the MGH Center for Skeletal Research NIH/NIAMS P30AR075042 (JAL), an Innovator Award from The Gillian Reny Stepping Strong Center for Trauma Innovation (JAL), a Brigham Research Institute microgrant (VK), and funds from BWH Department of Orthopedic Surgery (JAL).

