## supplemental material for "LGR6 is necessary for attaining peak bone mass and regulates osteogenesis through differential ligand use"

**SUPPLEMENTAL FIGURE LEGENDS**

**Supplemental Figure 1:**

Validation of Lgr6 siRNA-mediated knockdown in MSCs. (a) qPCR-derived RNA expression of Lgr6 72 hours post-transfection with 50nM or 100nM Lgr6 siRNA and control (siCon); data are analyzed by Student's t-test; ***p < 0.001. Error is reported as standard deviation. (b) Protein isolated from siRNA knockdown MSCs analyzed by western blot with anti-LGR6 or anti-GAPDH (control) antibodies. Reduced LGR6 protein levels found with 100nM Lgr6 siRNA. Each experiment was performed in technical duplicate and biological triplicate.

**Supplemental Figure 2:**

Knockdown of Lgr6 results in reduced osteoblast proliferation and differentiation in primary mouse calvarial osteoblasts (MCOs). (a) Cell proliferation quantified by WST-1 assay in Lgr6 knockdown (siLgr6) and control knockdown MCOs. (b and c) Relative RNA expression of Runx2 and Sp7 in Lgr6 knockdown MCOs relative to controls following 48 hours of osetoinduction. (d) Alkaline phosphatase (ALP) activity in knockdown MCOs 3 days post-osteoinduction. (e) Quantification of alizarin-red-S (ARS) staining of knockdown MCOs 21 days post-osteoinduction. Each experiment was performed in triplicate and analyzed by Student's t-test, *p < 0.05, **p < 0.01, ***p < 0.001. Error is reported as standard deviation.

**Supplemental Figure 3:** Generation of Lgr6 knockout (Lgr6-KO) mouse allele. (a) Schematic of wildtype and targeted Lgr6 genomic loci; black arrow represents direction of transcription starting from exon 1. Lgr6-KO allele has a deletion of exons 15 and 16 without the introduction of additional sequences. (b) PCR genotyping gels using genomic DNA from homozygous wildtype (+/+), heterozygous (+/-), and homozygous knockout (-/-) mice, for the wildtype allele (top) and the knockout allele (bottom); genomic position of primers WT.F/R and KO.F/R are shown in red in (a). (c) Lgr6-KO allele validation; Lgr6-WT and Lgr6-KO femoral bone marrow derived cDNA qPCR for Lgr6. Lgr6 Taqman assay Mm01291346 validates Lgr6-KO loss of expression ($p=0.004$).

**Supplemental Figure 4:**

siRNA mediated knockdown of Lgr6 in C3H10T1/2 cells does not affect adipogenesis or chondrogenesis. (a and a') Oil Red O staining of Lgr6 knockdown (a') and control (a) cells 12 days post-adipoinduction. (b) Quantification of Oil Red O staining shows no significant difference in adipodifferentiation. (c and c') Alcian Blue staining of Lgr6 knockdown (c') and control (c) cells 12 days post-chondroinduction. (d) Quantification of Alcian Blue staining shows no significant difference in chondrodifferentiation. Each experiment was performed in technical duplicate and biological triplicate and analyzed by Student's t-test; ns = not significant. Error is reported as standard deviation.

**Supplemental Figure 5:**

Validation of Lgr6 plasmid overexpression in MSCs. (a) qPCR-derived RNA expression of Lgr6 24 hours post-transfection with 25-100ng/ml of Lgr6 over-expression plasmid (pLgr6) or control (pCAG); data are analyzed by Student's t-test; *$p < 0.05$, **$p < 0.01$, ***$p < 0.001$. Error is reported as standard deviation. (b) Protein isolated from overexpression cells analyzed by western blot with anti-LGR6 or anti-GAPDH (control) antibodies. Increased expression of LGR6 is found with 50-100ng/ml pLgr6. Each experiment was performed in triplicate.

**Supplemental Figure 6:**

Lgr6 overexpression stimulates osteoblastogenesis in MCOs. (a) Cell proliferation quantified by WST-1 assay in Lgr6 overexpression (pLgr6) and control (pCAG) MCOs. (b) Alkaline phosphatase (ALP) activity 3 days post-osteoinduction and (c) Alizarin Red S staining (ARS) 21 days post-osteoinduction. Each experiment was performed in triplicate and analyzed by Student's t-test, *$p < 0.05$. Error is reported as standard deviation.

**Supplemental Fig 7:**

Lgr6 knockout mice have normal cortical bone parameters. MicroCT analysis of femurs from 8-week-old Lgr6 wildtype (WT) and Lgr6 knockout (KO) male and female mice. (a) Representative 3D images of cortical section from each sex and genotype. (b and c) No significant differences in (b) porosity, (c) cortical thickness and (d) tissue mineral density were found between genotypes. Cohort sizes: 14 female Lgr6-WT mice, 11 female Lgr6-KO mice, 11 male Lgr6-WT mice, and 12 male Lgr6-KO mice. Quantitative bone parameters were analyzed for significance by Student's t-test; sexes were analyzed separately, ns = not significant. Error is reported as standard deviation.

### TABLES

**Supplemental Table 1**

List of primers used in qPCR experiments, genotyping of Lgr6 knockout mouse allele, and putative CRISPR-Cas9 off-site target cut site PCR amplification and sequencing.

| **qPCR target gene** | **Primer 1 (5' to 3')** | **Primer 2 (5' to 3')** |
|---|---|---|
| HPRT | GGACTAATTATGGACAGGACTG | GCTCTTCAGTCTGATAAAATCTAC |

| | | |
|---|---|---|
| ALP | CGGATCCTGACCAAAAACC | TCATGATGTCCGTGGTCAAT |
| OPN | GGAAACCAGCCAAGGTAAGC | TGCCAATCTCATGGTCGTAG |
| Sp7 | CAAGAGTGAGCTGGCCTGA | TGGAGCCATAGTGAGCTTCTT |
| Runx2 | TCCACAAGGACAGAGTCAGATTAC | TGGCTCAGATAGGAGGGGTA |
| **Lgr6 allele genotyping** | **Primer 1 (5' to 3')** | **Primer 2 (5' to 3')** |
| Lgr6-WT | CATGGTCATTGTGTTAGGCTGA | TAGAGAGGAGATGGAGGTGAGA |
| Lgr6-KO | ACCCCTTGAACTCTCTTCCAAG | GCTTCTGACCTGTTTCCATTCA |
| **CRISPR allele generation off-target cutsite verification** | **Primer 1 (5' to 3')** | **Primer 2 (5' to 3')** |
| 1:134284169 | CCGGGCAAAGCAAGACATTC | AACAGGCTCTTCCTGGTTGG |
| 1:184695949 | GACTCTCTTGCCCTTCGCTG | CAGACCCCCAAAAGGACTCA |
| 8:13791972 | CACTCTCCTGGAAGCACGTT | AACGAGCAGAACAGACCCAG |
| 5:115576789 | CGATTGTGCTGTTTCCTGGC | AGCCACTCGCTGACAATGAA |
| 11:121068805 | GGGCATCATTGGCCTGAAAC | GAGGACAAGGTCCCAGACAC |
| 15:93503680 | TTAGCGAAACCTGCTCCCTG | TTCCATCACCCACAACCCAC |
| 1:153682542 | CCGAGCCACCTGAGAGTTTG | CTCTTCGGACCTCTTGGGAC |
| 1:152938331 | AGTGGACAGCAGAGGTCTCA | GGAATGCGCTGTGGGTGATA |
| 1:182312625 | CCACATGGCTCACAACCATC | CAGCCATGTCTGTGGCTACT |
| 1:55303945 | AGAGGACAAGACAAAGATGGAGAA | ACGACTTGTGAATCACCTTCCA |
| X:137571271 | CCTCGCACTCCTCTCTTTGAG | CCCACCAGGTTCACTACACT |
| 5:136982300 | CACAACCCTCCCATGACACA | CCCCGTATACCTGTGTGAGC |
| 7:112316435 | GGTCAGGCATCCTTCTAGAGC | TAAGGTGCTGGTATGGGAGGT |
| 11:116839590 | TCCACACTCGCTGAAGGAAA | CTGACTCTCGGGCATGAGG |
| 1:84216490 | AAATACAGGAAGCTGGCTTGAG | AGTTGCCCTCTGCATGTAAAAT |
| 1:14137156 | ACATGCTGGATGGTCTTTTCTT | CACTTTTGTGCACAGACAGACA |
| 1:19151281 | ACTCGTCCTAGATCCTCCCC | GCTTACAGGCAGAGTGTTTGC |
| 1:149045736 | TGTGGCACATATACCATTCAGC | GCCTCTCTGGAGAATTCCATTA |
| 1:62914362 | GCACACTGTAACCCTGAATGAA | GATGTGGGCTGCTCTAGTCTTT |
| 1:189193939 | GATGGACCATTCACATCACATC | CAAGCATGTGTTCTCCCACTAA |

### FIGURES

**Supplemental Figure 1**

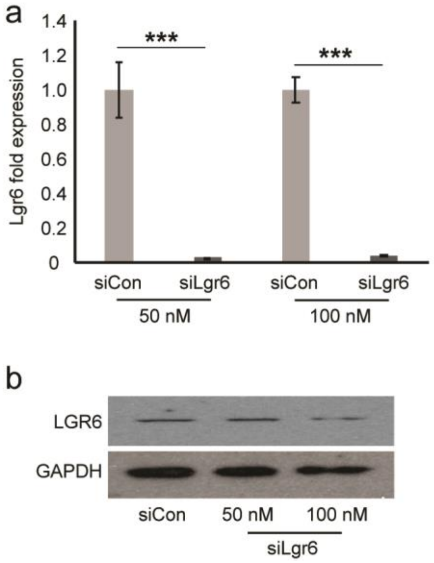

**Supplemental Figure 2**

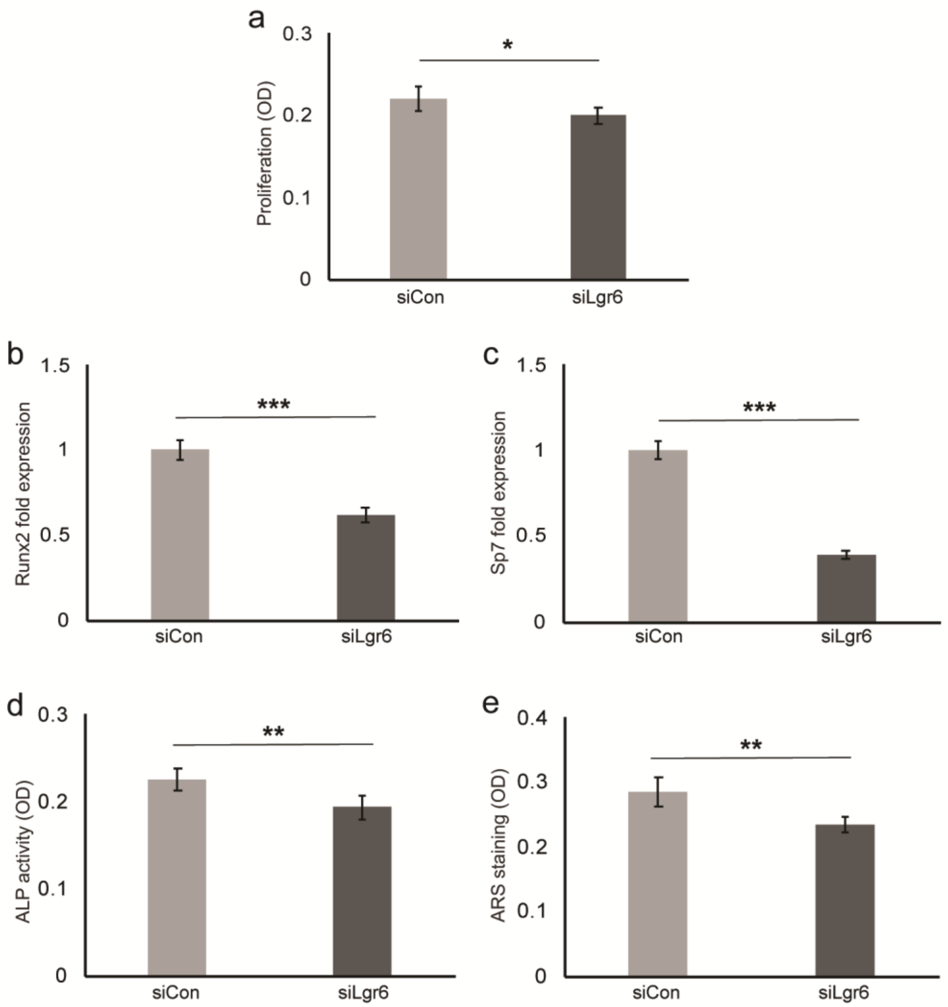

**Supplemental Figure 3**

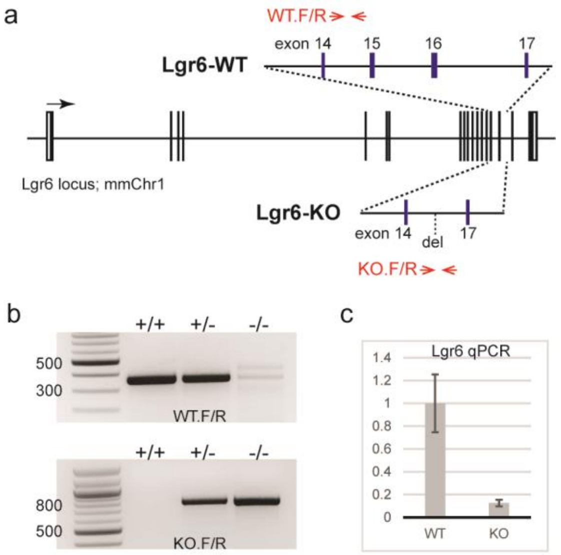

**Supplemental Figure 4**

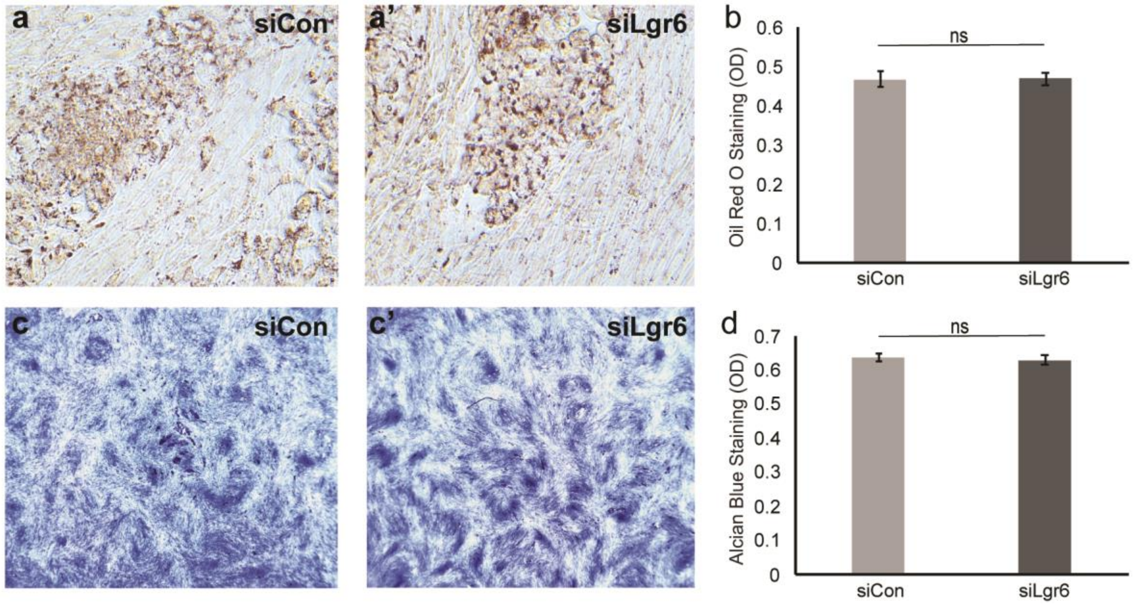

**Supplemental Figure 5**

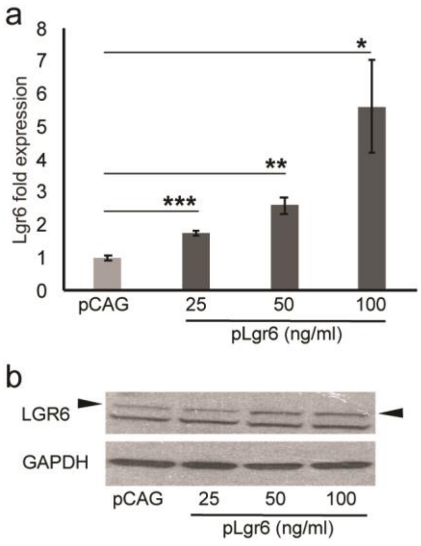

**Supplemental Figure 6**

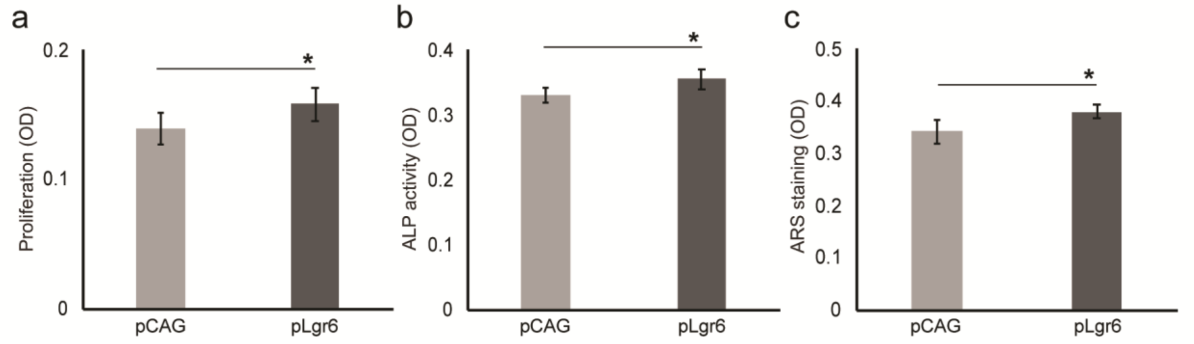

**Supplemental Figure 7**

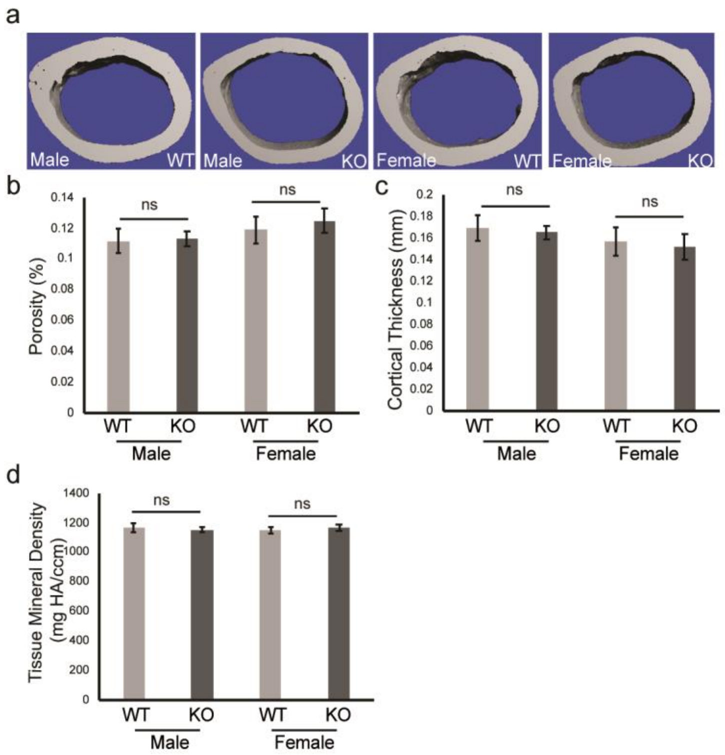
